# Aerosolized TLR Agonists Suppress Acute Sendai Virus Lung Infection and Chronic Airway Disease in Mice

**DOI:** 10.1101/668368

**Authors:** David L. Goldblatt, Jose R. Flores, Gabriella Valverde Ha, Ana M. Jaramillo, Sofya Tkachman, Carson T. Kirkpatrick, Shradha Wali, Belinda Hernandez, David E. Ost, Brenton L. Scott, Jichao Chen, Scott E. Evans, Michael J. Tuvim, Burton F. Dickey

## Abstract

Respiratory viral infections play central roles in the initiation, exacerbation and progression of asthma in humans. An acute paramyxoviral infection in mice can cause a chronic lung disease that resembles human asthma. We sought to determine whether reduction of Sendai virus lung burden in mice by stimulating innate immunity with aerosolized Toll-like receptor (TLR) agonists could attenuate the severity of chronic asthma-like lung disease. Treatment with 1 µM oligodeoxynucleotide (ODN) M362, an agonist of the TLR9 homodimer, and 4 µM Pam2CSK4 (Pam2), an agonist of the TLR2/6 heterodimer, within a few days before or after Sendai virus challenge, resulted in a ∼75% reduction in lung Sendai virus burden five days after challenge. The reduction in acute lung virus burden was associated with marked reductions 49 days after viral challenge in eosinophilic and lymphocytic lung inflammation, airway mucous metaplasia, lumenal mucus occlusion, and hyperresponsiveness to methacholine. Mechanistically, ODN/Pam2 treatment attenuated the chronic asthma phenotype by suppressing IL-33 production by type 2 pneumocytes, both by reducing the severity of acute infection and by downregulating Type 2 (allergic) inflammation. These data suggest that treatment of susceptible human hosts with aerosolized ODN and Pam2 at the time of a respiratory viral infection might attenuate the severity of the acute infection and reduce progression of asthma.

**One Sentence Summary:** Respiratory viral infections can induce chronic airway disease, and we find that stimulating innate immunity within the lungs of mice reduces the severity of acute infection and development of a chronic asthma phenotype.

## Introduction

Respiratory viral infections play a major role in the pathogenesis of asthma at every stage of disease – initiation, exacerbation, and progression (1-5). Regarding asthma initiation, children who experience a respiratory infection with rhinovirus or respiratory syncytial virus have a substantially increased chance of subsequently developing asthma. Regarding asthma exacerbations, new molecular techniques have allowed the identification of viral infections in more than 80% of exacerbation episodes in both children and adults. Last, progression of underlying asthma severity, defined as poorly reversible airflow obstruction and persistent symptoms, is associated with the frequency of respiratory viral infections. Rhinoviruses are the most common cause of asthma exacerbations, but paramyxoviruses such as respiratory syncytial virus and parainfluenza virus often cause severe disease and have a prominent role in disease initiation (1, 5).

While the epidemiologic association between respiratory virus infection and asthma is strong, a mechanistic explanation of the interaction has been lacking until recently. It now appears that the interaction operates in both directions. The presence of asthma results in an increased susceptibility to lower respiratory tract infection by viruses, possibly due at least in part to deficient interferon responses resulting from Type 2 immune deviation (2-4, 6, 7). Conversely, respiratory viral infection can result in long-term reprogramming of the lung immune environment towards a Type 2 polarity that results in parenchymal changes characteristic of asthma and COPD (2-4, 7, 8). The greatest source of mechanistic insight in this regard has come from studies of a mouse paramyxoviral lung infection with Sendai virus (SeV) by Holtzman and colleagues (9, 10). Analysis of this model has revealed that an acute SeV infection, in which virus is cleared within 12 days, can lead to a chronic, possibly lifetime, asthmatic phenotype. In susceptible mouse strains, the lungs 49 days after infectious challenge are characterized by Type 2 inflammation, epithelial mucous metaplasia, and hyperresponsiveness to bronchoconstrictor stimuli such as methacholine (6, 8-10). In recent work, this group has shown that the acute SeV infection leads to long-term elevated expression of IL-33 by lung epithelial cells that drives the persistent elaboration of high levels of IL-13 by immune cells within the lungs, which in turn induces the lung parenchymal changes of epithelial mucous metaplasia and smooth muscle hyperresponsiveness (11).

In view of the high incidence and substantial morbidity resulting from virus-induced asthma, treatments to mitigate the severity of respiratory virus infection are greatly needed. We have previously identified a pharmacologic means to stimulate innate immunity of lung epithelium that has shown activity against multiple bacterial, fungal, and viral pathogens in mice (12-18) and guinea pigs (19). The stimulus consists of a class C oligodeoxynucleotide that is an agonist of the TLR9 homodimer and Pam2CSK4 that is an agonist of the TLR2/6 heterodimer. Administration of these two agonists (hereafter called O/P) in a fixed molar ratio shows synergistic activity against microbial pathogens that is dependent upon MyD88 expression in lung epithelial cells and the generation of microbicidal reactive oxygen species (12, 14, 20, 21). We hypothesized that treatment with O/P at the time of respiratory infection with SeV would reduce lung virus burden during the acute infection and thereby attenuate the development of a late asthma phenotype.

## Results

### O/P treatment before SeV challenge reduces lung virus burden

The experimental paradigm is illustrated in Figure 1A. To test the first component of our hypothesis, mice were treated with aerosolized O/P one day before challenge with SeV because previous studies showed this is the interval of maximal benefit of a single O/P treatment for challenge with influenza virus or any of several bacterial pathogens (12-14). Lung SeV load was measured by RT-qPCR 5 days after challenge based upon this being the time of maximal viral burden in prior studies by others (9, 10, 22, 23) and our confirmatory study (fig. S1). Treatment with O/P resulted in a 70-80% reduction in SeV burden (Fig. 1B and fig. S1). Pilot studies titrating SeV challenge to mouse survival showed a steep dose-response relationship, with a 1.6-fold increase in virus load resulting in a decrease in mouse survival from 80% to 20% (Fig. 1C). Based on this tight relationship between viral burden and outcome, we anticipated a substantial change in late asthma phenotype with treatment, so proceeded to test the second component of the central hypothesis.

**Fig. 1.**
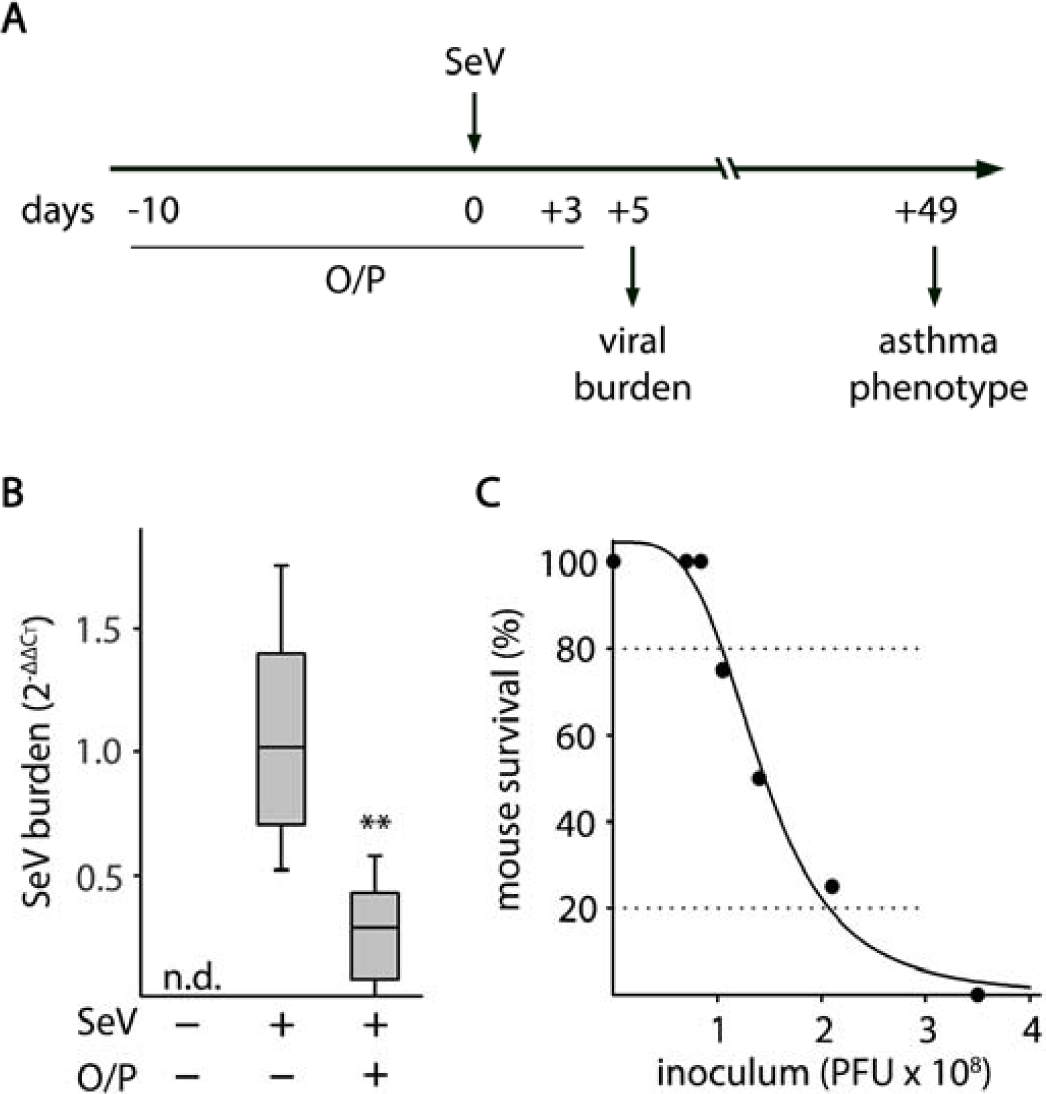
Experimental paradigm, effect of O/P treatment on lung SeV burden, and correlation between SeV burden and mouse survival. (**A**) Illustration of the experimental paradigm with O/P administered by aerosol a few days before or after SeV challenge, lung SeV burden assessed 5 days after challenge, and development of an asthma phenotype assessed 49 days after challenge. (**B**) Lung SeV burden measured by qRT-PCR 5 days after mice were challenged with SeV, with or without treatment one day earlier with O/P. (Boxes show median and interquartile range, whiskers show 10^th^ and 90^th^ percentiles; ^**^*P*<0.01 for [SeV^+^,O/P^+^] *vs* [SeV^+^,O/P^−^] by Student’s *t*-test; N=24 mice/group pooled from 4 experiments.) (**C**) Various SeV inocula were administered to determine the correlation between lung SeV burden and mouse survival. (N=4 mice/group in a representative experiment performed 5 times).

### O/P treatment before SeV challenge attenuates late asthma phenotype

Mice were treated with aerosolized O/P one day before challenge with SeV, and the severity of an asthma phenotype 49 days after viral challenge was measured. Treatment resulted in an 88% reduction in lung lavage eosinophil number, indicating the attenuation of persistent Type 2 lung inflammation, and an 86% reduction in lymphocyte number, suggesting a possible reduction in Th2 cells that sustain Type 2 lung inflammation (Fig. 2A-B). We next examined whether the reduction in Type 2 inflammation resulted in reduced effects on lung parenchymal cells. There was a 52% reduction in mucous metaplasia measured as airway epithelial mucin content (Fig. 2C-D), and a 53% reduction in airway hyperresponsiveness (AHR) measured as lung resistance to inflation after 30 mg/ml methacholine (Fig. 2E-F). These results suggest a causal relationship between the reduction in virus load from O/P treatment and the chronic asthma phenotype, so further evidence was sought by examining dose-response and time-response relationships.

**Fig. 2.**
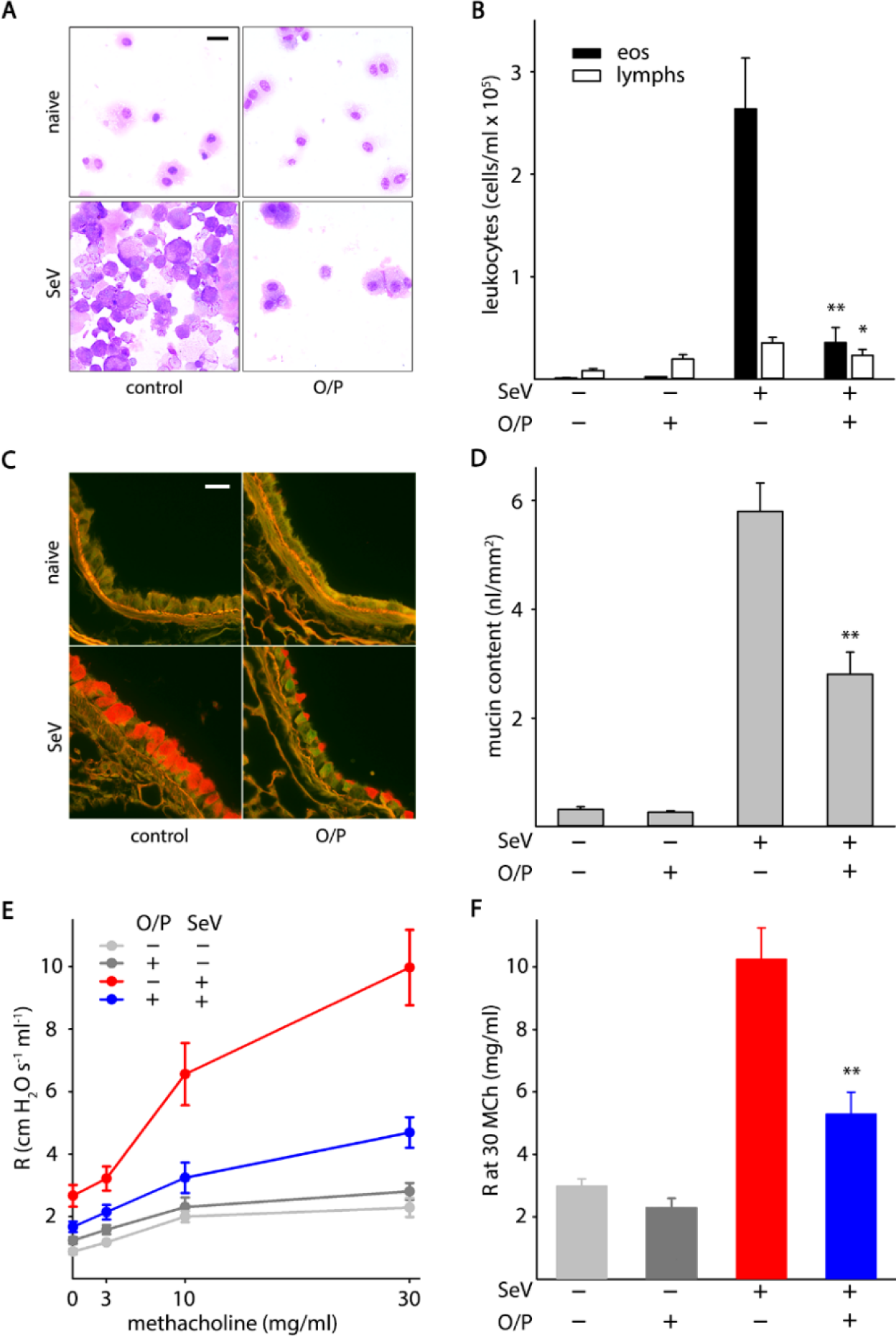
Effects of O/P treatment on asthma phenotype 49 days after SeV challenge. (**A**) Leukocytes obtained by lung lavage were pelleted onto glass slides by centrifugation and stained with Wright-Giemsa. Scale bar = 20 µm. (**B**) Eosinophils and lymphocytes in lung lavage fluid were enumerated. (Bars show mean ± SEM; ^*^*P*<0.05 and ^**^*P*<0.01 for [SeV^+^,O/P^+^] *vs* [SeV^+^,O/P^−^] by Student’s *t*-test; N=21 mice/group pooled from 3 experiments.) (**C**) Airway epithelium stained with PAFS to demonstrate intracellular mucin in red. Scale bar = 20 µm. (**D**) Quantification of intracellular mucin content by image analysis of airway as in (C). (Bars and *P* values as in (B); N=22-25 mice/group pooled from 5 experiments.). (**E**) Dose-response relationship between the concentration of aerosolized methacholine and total respiratory resistance measured by forced oscillation. (Points show mean ± SEM; N=9-14 mice per group pooled from 3 experiments.) (**F)** Plot of total respiratory resistance at 30 mg/ml methacholine (MCh) using data from E. (Bars and *P* values as in (B)).

### Parallel dose-response relationships between acute SeV burden and late asthma phenotype

Our previous studies had shown that the combination of 1 µM ODN and 4 µM Pam2CSK4 in 4 ml (1X O/P) is on the inflection of the upper plateau of the dose-response curves for airway epithelial activation as measured by cytokine release (15) and for antimicrobial resistance as measured by the survival of mice from a microbial challenge (15, 20). Now, we evaluated the responses to two halving doses of O/P below the 1X dose to interrogate the steep portion of the dose-response curve, and to two doubling doses above the 1X dose to confirm that responses were saturated. Mice treated one day before SeV challenge showed dose-dependent reductions in lung virus burden 5 days after challenge (Fig. 3A), and in lung lavage eosinophil and lymphocyte numbers (Fig. 3B) and mucous metaplasia (Fig. 3C) 49 days after challenge.

**Fig. 3.**
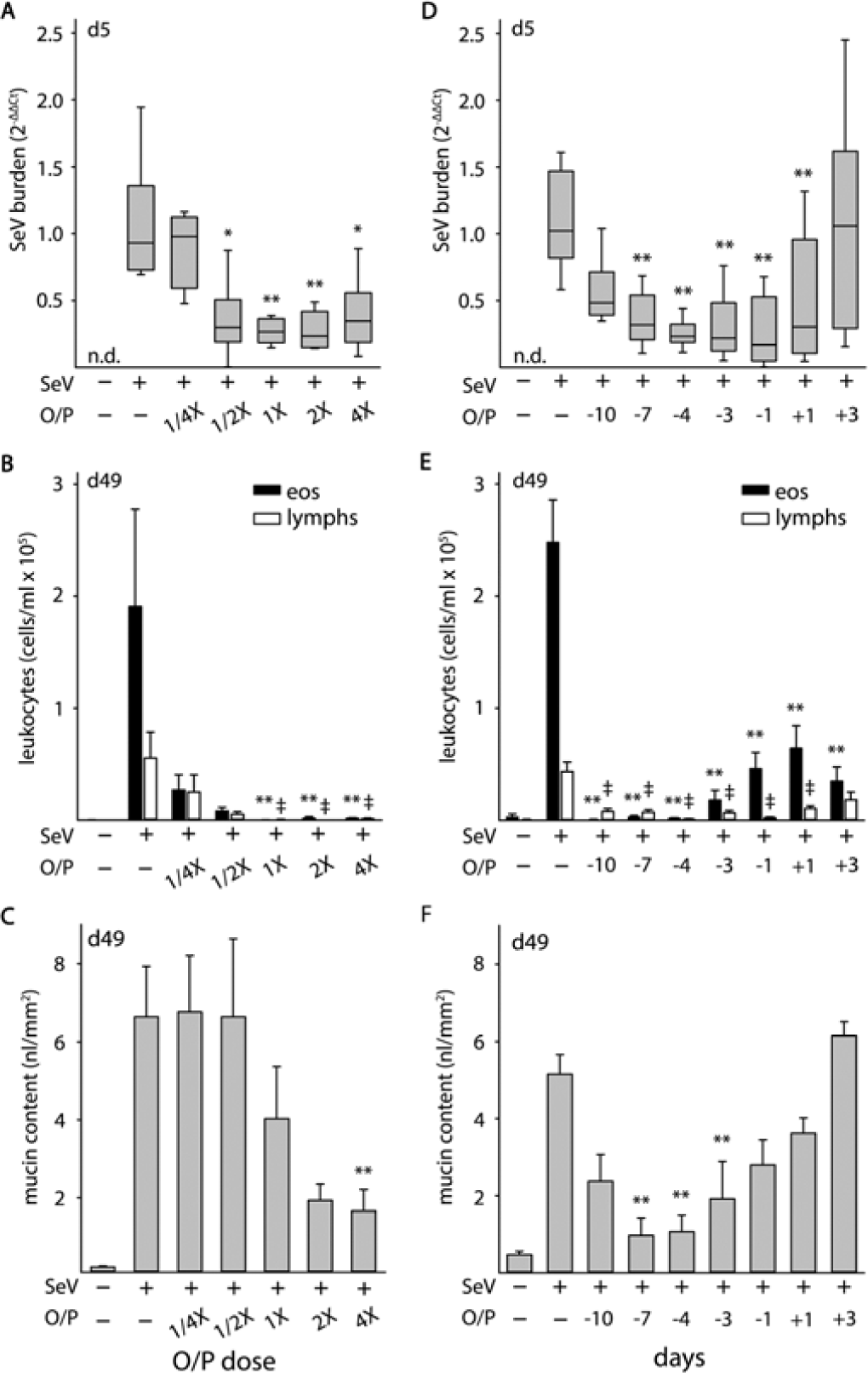
Dose and temporal response relationships between O/P treatment and acute lung SeV burden and late asthma phenotypes. (**A**) Relationship between O/P dose and lung SeV burden 5 days after SeV challenge. (Boxes show median and interquartile range, whiskers show 10^th^ and 90^th^ percentiles; ^*^*P*<0.05 and ^**^*P*<0.01 for ANOVA on ranks with Dunn’s test for multiple comparisons against [SeV^+^,O/P^−^] control; N=5-6 mice/group in a single experiment; n.d. = not detectable.) (**B**) Relationship between O/P dose and lung lavage eosinophil and lymphocyte numbers 49 days after SeV challenge. (Bars show mean ± SEM; ^**^*P*<0.01 as in (A), L*P*<0.01 for comparisons on lymphocytes; N=5 mice/group in a single experiment.) (**C**) Relationship between O/P dose and intracellular mucin content 49 days after SeV challenge. (Bars show mean ± SEM; ^**^*P*<0.01 as in (A); N=6-7 in a single experiment.) (**D**) Relationship of the interval in days between O/P treatment and SeV challenge in SeV lung burden 5 days after challenge. (Box plot and *P* values as in (A); N=25-39 mice/group pooled from 7 experiments.) (**E**) Relationship of the interval between O/P treatment and SeV challenge in lung lavage eosinophil and lymphocyte numbers 49 days after challenge. (Bars and *P* values as in (B); N=25-29 mice/group pooled from 6 experiments). (**F**) Relationship of the interval between O/P treatment and SeV challenge in intracellular mucin content 49 days after challenge. (Bars and *P* values as in (C); N=6-12 mice/group pooled from 3 experiments.)

The correlation among these dose-response relationships supports a mechanistic interaction between the reduction in acute viral lung burden by O/P treatment and the attenuation of a late asthma phenotype. However, it is notable that the maximal effect of O/P on lung virus burden appeared to be at the 1/2X dose, on lung lavage eosinophil and lymphocyte levels at the 1X dose, and on mucous metaplasia at the 2X dose. These small concentration-dependent differences among outcomes were consistent across multiple experiments, and suggest that small unmeasurable differences in lung virus burden at higher O/P concentrations (Fig. 3A) are sufficient to cause measurable differences in persistent lung inflammation (Fig. 3B), leading in turn to downstream effects on lung parenchymal phenotype (Fig. 3C). This interpretation is consistent with the steep dose-response relationship between lung virus burden and mouse mortality (Fig. 1C).

### Time-response relationships between acute SeV burden and late asthma phenotypes

To test temporal relationships, mice were treated with a 1X dose of O/P at varying intervals before and after challenge with SeV. Our previous studies showed variable durations of protection depending upon the intensity of the infectious challenge, with benefit observed with a pretreatment interval as long as eight days before a bacterial challenge of mild intensity (12), but as short as three days before a challenge of severe intensity (24). For treatments given after challenge, we have found a great difference between bacterial and viral pathogens, with a therapeutic window of less than one day for bacterial pathogens (12, 24), but of several days for influenza virus (13, 21). Based on these prior observations, responses to O/P from 10 days before to 3 days after SeV challenge were evaluated. A reduction in virus burden was seen from 10 days before challenge to 1 day after challenge (Fig. 3D). Reductions in eosinophils and lymphocytes were observed at all time points (Fig. 3E), and for mucous metaplasia at all time points except 3 days after challenge (Fig. 3F). Overall, there was a trend towards greater effects with shorter intervals between treatment and challenge. Similar to the dose-response relationship, the time-dependent effects of O/P treatment on leukocyte numbers were greater than on SeV burden (Fig. 3E-F).

The efficacy of O/P given 10 days before SeV challenge in these studies was surprising. To test whether O/P might be influencing the development of a chronic asthma phenotype in the SeV infection model by exerting an immunomodulatory effect, O/P was administered at time points substantially outside its known window of antimicrobial efficacy. Treatment with O/P 20 days before SeV challenge resulted in a reduction 49 days after challenge in eosinophils by 85% and lymphocytes by 75%; similarly, treatment 20 days after SeV challenge resulted in a reduction 49 days after challenge in eosinophils by 73% and lymphocytes by 74% (fig. S2). These findings indicate that O/P treatment attenuates the late asthma phenotype in SeV-challenged mice by a combination of a reduction in acute lung virus burden and long-term immunomodulation away from a T2 lung inflammatory milieu.

### O/P treatment reduces airway mucus occlusion and acidophilic pneumonitis

We observed multiple yellow nodules on the surface of the lungs of untreated mice challenged with SeV 49 days earlier (Fig. 4A). Transillumination allowed us to enumerate nodules within the lungs (Fig. 4B), and to measure their mean diameter at 1.1 mm (SEM = 45 µm, N = 37). Mice treated with O/P one day before SeV challenge had a 92% reduction in nodule number (Fig. 4C). Histopathologic examination of the nodules showed alveolar accumulation of foamy macrophages, eosinophils, and Charcot-Leyden crystals (Fig. 4D-E), characteristic of “acidophilic pneumonitis” (25, 26). The identity of the macrophages was confirmed by immunofluorescence staining for CD68 (fig. S3D), and Sudan black staining showed the macrophages to be heavily laden with lipids (Fig. 4F), even more so than the normal staining of lamellar bodies in alveolar T2 cells (fig. S3E).

**Fig. 4.**
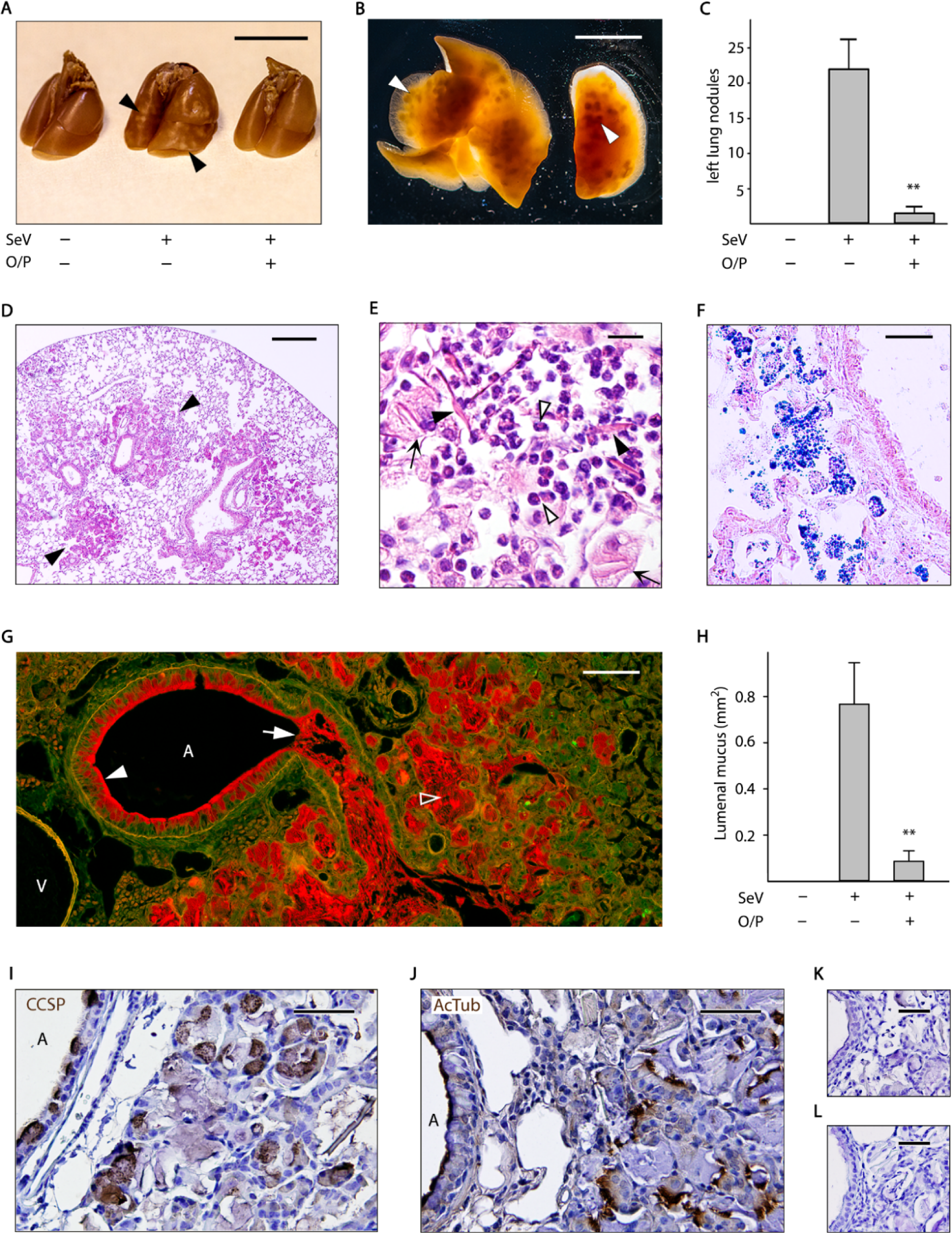
Airway mucus occlusion, acidophilic pneumonitis, and alveolar bronchiolization 49 days after SeV infection. (**A**) The lungs of mice treated or not with O/P, then challenged or not with SeV one day later, then sacrificed 49 days later with lungs inflated with 10% formalin to 20 cm H_2_O pressure. Arrowheads points to nodules on the lung surface. Scale bar = 1 cm. (**B**) Transilluminated lungs from a mouse challenged with SeV but not treated with O/P, as in (A).Arrowheads point to nodules in the lung interior. Scale bar = 1 cm. (**C**) Enumeration of nodules in the left lungs of mice treated with O/P and challenged with SeV as in (A). (Bars indicate mean ± SEM; ^**^*P*<0.01 for [SeV^+^,O/P^+^] *vs* [SeV^+^,O/P^−^] by Student’s *t*-test; N=4-5 in a single experiment.) (**D**) Section of the lung of a mouse challenged with SeV but not treated with O/P, stained with H&E. Arrowheads point to nodules. Scale bar = 0.5 mm. (**E**) High magnification image of a nodule from a mouse as in (D), with closed arrowheads pointing to Charcot-Leyden crystals, open arrowheads pointing to eosinophils, and arrows pointing to foamy macrophages. Scale bar = 30 µm. (**F**) Image of a nodule from a mouse as in (D), but with lungs frozen and stained with Sudan black to show lipids within macrophages. Scale bar = 50 µm. (**G**) Section of the lung of a mouse challenged with SeV but not treated with O/P, fixed by immersion in methacarn and stained with PAFS. Closed arrowhead points to abundant intracellular mucin, arrow points to lumenal mucus occlusion, open arrowhead points to mucus in the alveolar region. A = airway, V = vessel. Scale bar = 100 µm. (**H**) The area of lumenal mucus of mice treated or not with O/P, then challenged or not with SeV, then fixed and stained as in (G), with left lungs sectioned at fixed intervals using a precision cutting tool. (Bars indicate mean ± SEM; ^**^*P*<0.01 for [SeV^+^,O/P^+^] *vs* [SeV^+^,O/P^−^] by Mann-Whitney *U* test; N=13-15 mice/group in a single experiment.) (**I-L**) Alveolar region of a mouse challenged with SeV but not treated with O/P, stained with antibodies against (I) CCSP or (J) acetylated tubulin (both brown), and counterstained with H&E (blue). A = airway lumens. Primary antibodies against (K) CCSP or (L) acetylated tubulin were omitted to assess the specificity of antibody staining. Scale bar in all 4 images = 50 µm.

We hypothesized that mucus plugs caused airway obstruction leading to lipid-laden macrophage accumulation in alveoli. To evaluate this, uninflated lungs were fixed with methacarn to minimize changes in mucus volume and position, and then stained with PAFS to visualize mucins (27). Supporting this hypothesis, widespread mucus plugs were visible in the airways of mice challenged 49 days earlier with SeV (Fig. 4G, arrow), and were greatly reduced in the airways of mice treated with O/P one day before challenge (Fig. 4H). The mucus plugs were contiguous with areas of intense airway epithelial mucous metaplasia (Fig. 4G, filled arrowhead), and the plugs contained both Muc5ac and Muc5b (fig. S3A-B). In the process of measuring airway lumenal mucus using a precision lung-slicing tool (fig. S4), we observed that lungs from untreated SeV-challenged mice were larger (25-30% greater rostral-caudal length) than those from unchallenged or treated and challenged mice, consistent with the obstructive hyperinflation observed at autopsy in the lungs of human subjects who die from asthma (28).

Red PAFS staining also extended into alveolar regions distal to mucus plugs (Fig. 4H, open arrowhead), and these areas were also positive for Muc5ac and Muc5b immunohistochemical staining (fig. S3A), with relatively more Muc5ac staining in airways and relatively more Muc5b staining in alveolar regions. To assess whether mucus in alveolar regions might be due to local mucin production resulting from bronchiolization of alveolar epithelium, we stained for CCSP and acetylated tubulin, markers of airway secretory and ciliated cells, respectively. This showed normal staining in conducting airways as well as abnormal staining in alveolar regions (Fig. 4I-J).

### O/P treatment attenuates the disruption of epithelial mosaicism

We considered that a possible additional contributor to airway lumenal mucus accumulation besides mucin hyperproduction might be impaired ciliary clearance because we and others have observed extensive epithelial cell death in the first 2 weeks after SeV challenge (21, 23, 29-31). The airway epithelium was examined 2-7 weeks after SeV challenge by whole mount immunofluorescence microscopy, and the normal mosaic of secretory and ciliated cells was altered in multiple ways. Secretory cells, marked by CCSP staining, became more prominent than ciliated cells, marked by acetylated tubulin staining (Fig. 5 and fig. S5). This prominence was due to a modest increase in secretory cell number (fig. S5B) and a more substantial increase in secretory cell size and intensity of CCSP staining (Fig. 5 and fig. S5A). The increase in secretory cell size is likely due to distension by intracellular mucin content (Fig. 2C and 4G, and fig. S3A-B and S5C), as we have shown previously during mucous metaplasia (32, 33). The increase in intensity of CCSP staining (Fig. 5 and fig. S5A) likely reflects a combination of increased expression and redistribution from small apical secretory granules in naïve cells into large mucin-containing granules during Type 2 inflammation (32, 33). Besides these changes in cell proportions and size, an alteration of the normal epithelial polygonal mosaic was revealed by staining cell boundaries using antibodies to E-cadherin (Fig. 5), with cells taking on more elongated shapes and occurring in abnormally large clusters of secretory cells. We observed these changes as early as 2 weeks after SeV challenge (fig. S5), found they partially persisted partially through 7 weeks (not shown), and that there was a trend towards improvement by pretreatment with O/P (fig. S5B).

**Fig. 5.**
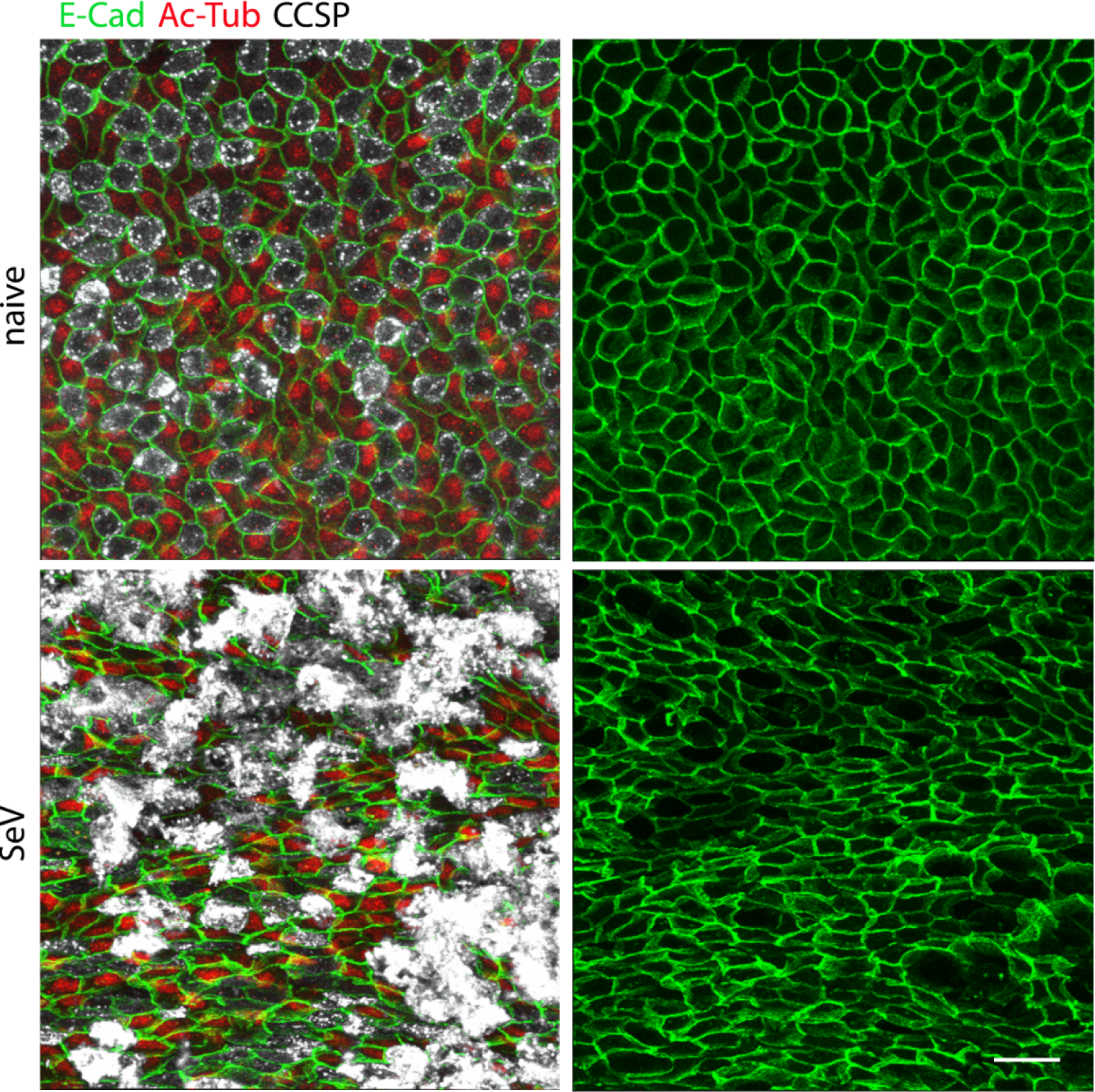
SeV infection alters the normal airway epithelial mosaic. Whole mount immunofluorescence images of proximal portions of the axial bronchi of mice infected or not with SeV 5 weeks earlier. Images on the left show staining for E-cadherin (E-Cad, green) to outline cell borders, acetylated tubulin (Ac-Tub, red) to identify ciliated cells, and club cell secretory protein (CCSP, white) to identify secretory cells. Images on the right only show staining for E-cadherin to better illustrate changes in cell shape and pattern. Scale bar = 20 µm.

### O/P treatment before SeV challenge reduces lung IL-33 expression 49 days later

Holtzman and colleagues have shown that the effects of acute SeV infection are translated into a chronic asthma phenotype by persistently increased IL-33 expression in the lungs (11). Therefore we measured the effect of O/P treatment during acute SeV infection on IL-33 expression 49 days later. Naïve mice that were neither challenged with SeV nor treated with O/P showed scattered faint immunohistochemical staining for IL-33 in cells with the localization (corners of polygonal alveoli) and morphology (cuboidal and protruding into the alveolar lumen) of type 2 pneumocytes (Fig. 6A). This finding is consistent with reports of substantial expression of IL-33 in type 2 pneumocytes of healthy mice (34, 35). There was no apparent change in IL-33 expression in the lungs of unchallenged mice treated with O/P, but mice challenged with SeV and not treated with O/P showed a dramatic increase in the intensity and frequency of IL-33 staining of type 2 pneumocytes and faint staining of macrophages (Fig. 6A). This was quantified by immunofluorescence staining (Fig. 6B), and mice treated with O/P and then challenged one day later with SeV showed a marked reduction in IL-33 staining (Fig. 6A-B).

**Fig. 6.**
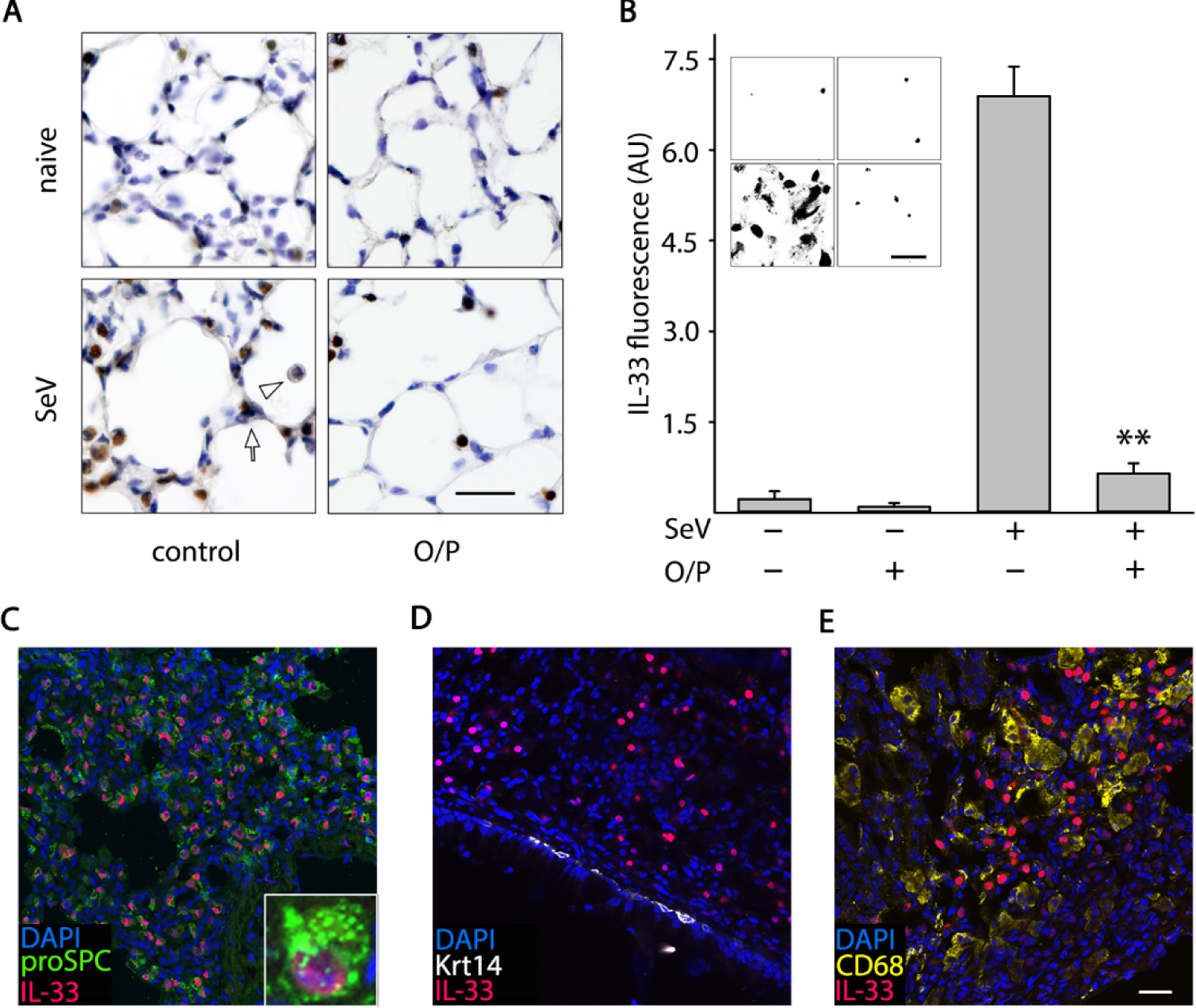
Persistent IL-33 expression after SeV challenge. (**A**) Images of the lungs of mice treated or not with O/P, then challenged or not with SeV one day later, then sacrificed 49 days later. Brown color is immunohistochemical staining for IL-33. Arrow points to an intensely stained alveolar T2 cell, arrowhead points to a faintly stained macrophage. Scale bar = 100 µm.(**B**) Same groups as in (A), but with fluorescence staining for IL-33. Inset shows masking for staining greater than background. Scale bar = 100 µm. (Bars indicate mean ± SEM; ^**^*P*<0.01 for [SeV^+^,O/P^+^] *vs* [SeV^+^,O/P^−^] by Student’s *t*-test; N=3 mice/group in a single experiment, with 3 fields examined per mouse.) (**C**) Fluorescence staining for proSPC to identify alveolar T2 cells (green), IL-33 (red), and DAPI to identify nuclei (blue). Inset shows IL-33 expression in a T2 cell. (**D**) Fluorescence staining for cytokeratin 14 (Krt14) to identify airway basal cells (white), and IL-33 and DAPI as in (C), shows no apparent expression of IL-33 in basal cells. (**E**) Fluorescence staining for CD68 to identify macrophages (yellow), and IL-33 and DAPI as in (C), shows no apparent expression of IL-33 in macrophages. Scale bar for (C, D, E) = 200 µm, and for inset in (C) = 30 µm.

Immunofluorescence co-localization of IL-33 staining with lineage markers showed that high-level IL-33 expression in SeV-challenged mice occurred exclusively in type 2 pneumocytes, with 97% of IL-33-positive cells also positive for pro-SP-C (Fig. 6C) and no measurable IL-33 staining in Krt14-positive basal cells of the conducting airways (Fig. 6D) or in CD68-positive alveolar macrophages (Fig. 6E) (n = 100 cells in 3 mice for each lineage marker). In SeV-challenged mice, 71% of pro-SP-C-positive cells were positive for IL-33 staining (n = 100 cells in 3 mice).

### Multiple doses of O/P neither induce tachyphylaxis nor augment efficacy

We next sought to determine whether multiple O/P doses induce tachyphylaxis or improve outcomes. Repetitive treatment with 2 to 4 doses of O/P given daily or separated by 2 or 3 days between doses, with the final dose for each regimen given one day before SeV challenge (Fig. 7A), did not result in any measurable differences in the reduction of lung viral burden by O/P compared to a single treatment (Fig. 7B). Therefore we concluded that there is no tachyphylaxis of the antiviral effect of O/P treatment, similar to what we have previously observed with influenza virus and bacterial pathogens (36, 37), so we proceeded to test whether multiple dosing improved outcomes.

**Fig. 7.**
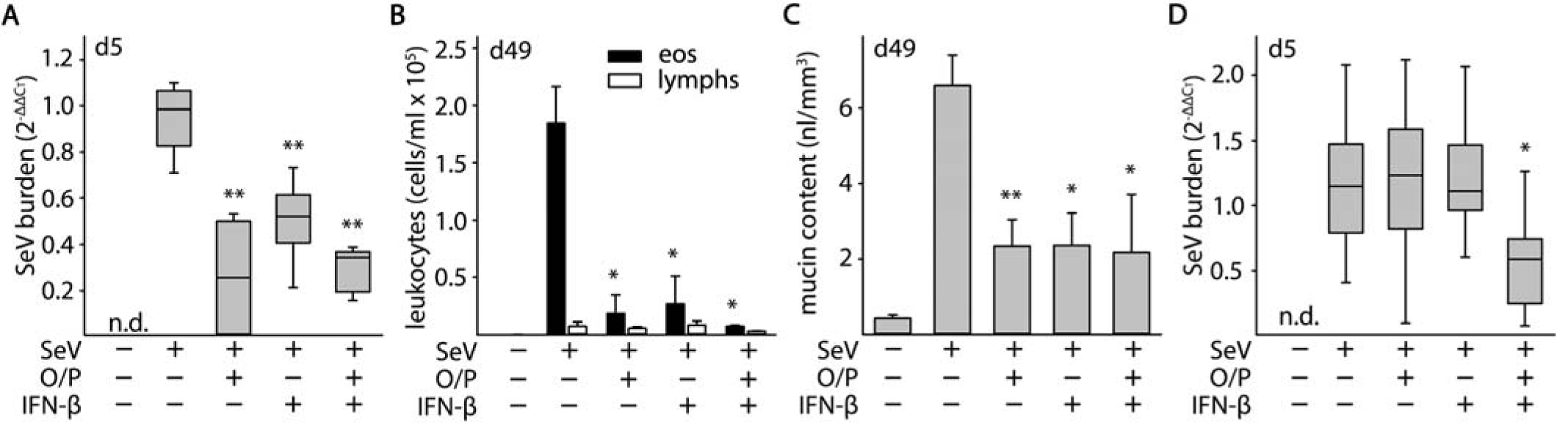
Effects of multiple O/P doses on tachyphylaxis and efficacy. (**A**) Experimental paradigm illustrating the timing of O/P doses for each experimental group in black triangles in relation to the SeV challenge given on day 0. (**B**) Lung SeV burden by qRT-PCR 5 days after mice were challenged with SeV, with or without pretreatment with O/P as outlined in (A). (Boxes show median and interquartile range, whiskers show 10^th^ and 90^th^ percentiles; ^**^*P*<0.01 for analysis by ANOVA on ranks with Dunn’s test for multiple comparisons against [SeV^+^,O/P^−^] control; N=4-7 mice/group in a single experiment; n.d. = not detectable.) (**C**) Lung SeV burden by RT-qPCR 5 days after mice were challenged with SeV, with or without treatment with O/P one day before challenge (-1), one day after challenge (+1), or both (-/+1). (Box plot and *P* values as in (B); bar and labeled *P* values for single comparison between [SeV^+^,O/P^−1^] *vs* [SeV^+^,O/P^−/+1^] by Student’s *t*-test; N=14-25 mice/group pooled from 3 experiments.) (**D**) Eosinophils and lymphocytes in lung lavage fluid were enumerated 49 days after SeV challenge, with or without treatment with O/P as in (C). (Bars show mean ± SEM; ^*^*P*<0.05 and ^**^*P*<0.01 as in (B), ^**†**^*P*<0.05 and ^□^ *P*<0.01 for comparisons on lymphocytes; bar and labeled *P* values as in (C); N=8-11 mice/group pooled from 2 experiments.) (**E**) Quantification of intracellular mucin content of airway epithelium 49 days after SeV challenge, with or without treatment with O/P as in (C). (Bars and *P* values as in (C); N=8-11 mice/group pooled from 2 experiments.)

Treatment with O/P both one day before and one day after SeV challenge did not reduce lung virus burden more than a single treatment before challenge (Fig. 7C). Similarly, when late asthma outcomes were examined, treatment both before and after challenge was no more effective than a single treatment before challenge in reducing lung lavage eosinophils and lymphocytes (Fig. 7D) or epithelial mucin content (Fig. 7E). In each case, treatment one day after challenge was less effective than treatment one day after challenge (Fig. 7C-E).

### Comparison of O/P to IFN-β effects on lung SeV burden and asthma phenotype

IFN-β has been shown to be a key signaling molecule in defense against respiratory viral infections, and aerosolized IFN-β has progressed to clinical trials for the treatment of virus-induced asthma exacerbations (38). To compare treatment with O/P to IFN-β, we first developed an IFN-β treatment paradigm in mice. Serum CXCL10 levels were used as a biomarker of anti-viral activity in the published clinical trial (38), so we measured CXCL10 levels in BALF in mouse pilot studies. The time of peak CXCL10 level in BALF was 2 h after aerosolized IFN-β treatment (fig. S6A), and a dose of 400,000 IFN-β units resulted in a CXCL10 level that lay on the edge of the upper plateau of the dose-response curve (fig. S6B), so this treatment timing and dose was used in efficacy studies. Reductions in lung SeV burden acutely (Fig. 8A), and in lavage eosinophils and lymphocytes (Fig. 8B) and in epithelial mucin content (Fig. 8C) 49 days after SeV challenge, were similar between O/P and IFN-β treatments.

**Fig. 8.**
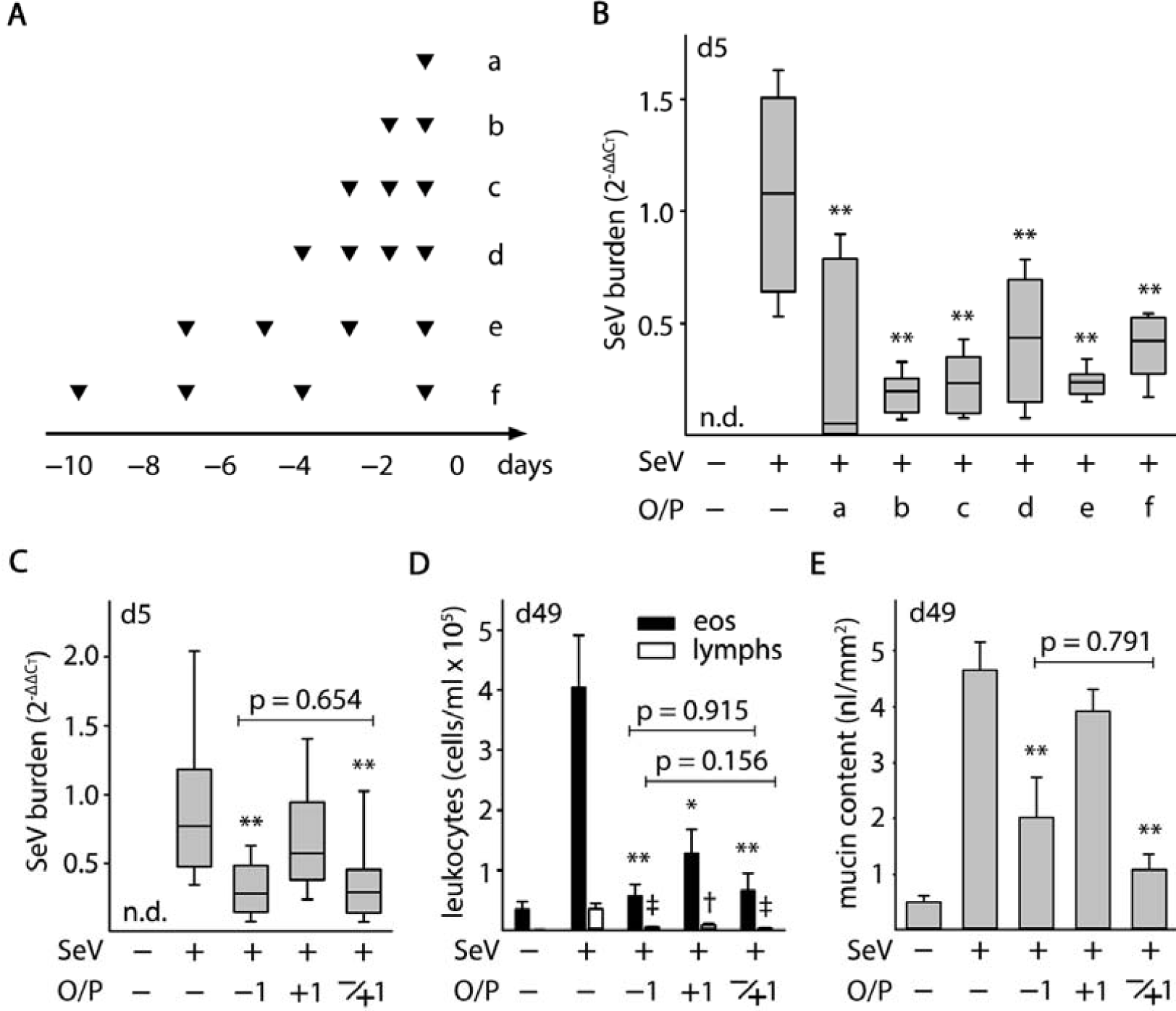
Comparison of the effects of O/P and IFN-β on acute lung SeV burden and late asthma phenotypes. (**A**) Lung SeV burden by qRT-PCR 5 days after mice were challenged with SeV, with or without pretreatment one day earlier with O/P, IFN-β, or both. (Boxes show median and interquartile range, whiskers show 10^th^ and 90^th^ percentiles; ^*^*P*<0.05 and ^**^*P*<0.01 for [SeV^+^,drug^+^] *vs* [SeV^+^,drug^−^], with *P*>0.05 for all other possible pairwise comparisons by one-way ANOVA with Holm-Sidak’s test for multiple comparisons; N=4-6 mice/group in a single experiment; n.d. = not detectable.) (**B**) Eosinophils and lymphocytes in lung lavage fluid were enumerated 49 days after SeV challenge as in (A). (Bars show mean ± SEM; *P* values as in (A); N=4-6 mice/group in a single experiment.) (**C**) Quantification of intracellular mucin content of airway epithelium 49 days after SeV challenge as in (A). (Bars and *P* values are as in (A); N=3-4 in a single experiment). (**D**) Lung SeV burden by RT-qPCR 5 days after mice were challenged with a high dose of SeV (2.5 × LD_50_), with or without pretreatment one day earlier with O/P, IFN-β, or both. (Boxes show median and interquartile range, whiskers show 10^th^ and 90^th^ percentiles, n.d.=not detectable; ^*^*P*<0.05 for both drugs together *vs* either drug alone by one-way ANOVA with Holm-Sidak’s test for multiple comparisons; N=3 for the uninfected group, 10 for all other groups in a single experiment.)

O/P treatment does not cause a rise in Type I interferons (12-14, 21), so we wondered whether O/P and IFN-β might have an additive effect. Supporting separate mechanisms of action, pilot studies showed that treatment with O/P or IFN-β elicited different cytokine profiles, with O/P causing a substantially greater rise in CXCL2 and TNF-α, and IFN-β causing a greater rise in CXCL10 (fig. S6C). In addition, both treatments together caused a synergistic rise in IL-6 (fig. S6C). Despite these promising preliminary data, there was no apparent benefit to treating mice with both drugs simultaneously on lung SeV burden, lavage eosinophils and lymphocytes, or epithelial mucin content (Fig. 8A-C). To further test the possibility of additive activity of the two drugs, we gave a high-intensity SeV challenge (2.5 × LD_50_), for which there was no apparent reduction in lung SeV burden by either drug alone (Fig. 8D). However, the combination of both drugs reduced lung SeV burden significantly compared to the absence of treatment or to treatment with either drug along (Fig. 8D).

## Discussion

Recent advances in molecular diagnostics have strengthened the tight association between viral infection and asthma initiation, exacerbation, and progression (1-4). Complementing this data, recent work with animal models has supported a causal role of respiratory viral infections in promoting long-term asthma-like disease and has enabled the identification of critical pathways in translating acute infection into chronic disease (5, 6, 11, 19, 30, 39). Together, these developments highlight the importance of attenuating the severity of respiratory viral infections to prevent chronic airway diseases. We have previously identified a means of effectively stimulating the airway epithelium to induce innate immune resistance to respiratory viral infection (12-21). Here we show that treatment of mice with this efficacious combination of aerosolized TLR ligands around the time of SeV infection attenuates chronic lung disease, providing proof-of-principle that this intervention might have clinical efficacy.

Based upon our prior studies showing that treatment with O/P results in rapid pathogen killing within the lungs, we hypothesized that the major mechanism of action of O/P in ameliorating chronic SeV-induced lung disease would be by reducing lung virus burden. The close dose-response relationship between acute SeV lung burden and chronic asthma phenotypes supports such a causal relationship (Fig. 3A-C). The relationship between SeV lung burden and acute disease is steep, with a 1.6-fold decrease in viral burden resulting in a 4-fold decrease in mouse death (Fig. 1C). Similarly, reduction of SeV lung burden by 70-80% has a dramatic effect on chronic disease (Fig. 1B, 3A, 3D, 7B-C, 8A). This contrasts with the need for multiple log reductions in pathogen burden in chronic mycobacterial lung infection and acute bacterial blood infections to achieve comparable benefit, reflecting the capability of small numbers of residual organisms to perpetuate disease or develop drug resistance in those infections.

The major mechanism of pathogen killing with O/P treatment is the activation of reactive oxygen generators in airway epithelial cells, which persists at a high level for several days, then slowly declines to its baseline level (14, 18, 21). This is consistent with the reduction in SeV lung burden when mice are treated in the few days before or after challenge (Fig. 1D). However, the substantial reduction in lung inflammation 49 days after SeV challenge observed in mice treated 10 days before challenge with O/P (Fig. 3E), when there is not a significant effect on lung viral burden in this (Fig. 3D) or our previous studies (13, 21), suggested an additional effect besides viral killing. To further explore this, we treated mice with O/P either 20 days before or 20 days after SeV challenge and observed substantial effects on lung inflammation 49 days after SeV challenge (fig. S2). This suggests that O/P also exerts an immunomodulatory effect on lung epithelium that additionally attenuates the chronic inflammatory lung phenotype independent of its antiviral effect. Holtzman and colleagues have shown that persistent expression of high levels of IL-33 in the lungs of SeV-challenged mice connects the acute infection to the chronic disease (11). Consistent with this, O/P treatment markedly reduced IL-33 expression in alveolar T2 cells (Fig. 6), which are thought to signal to distal airways in an “outside-in” manner (34, 35, 40, 41). The degree to which the reduction in IL-33 expression by O/P treatment reflects decreased epithelial cell injury and death and the degree to which it reflects immune modulation will require further study.

A prominent feature of chronic lung disease in SeV-challenged mice was the presence of multiple millimeter-sized nodules (Fig. 4A-C). By microscopic analysis, these were comprised of areas of acidophilic pneumonitis (25, 26), characterized by lipid-laden foamy macrophages, eosinophilic and lymphocytic inflammation, and Charcot-Leyden crystals (Fig. 4D-F), as well as areas of bronchiolization characterized by alveolar regions replaced by nests of club and ciliated cells (Fig. 4I-J). Abundant mucus was observed occluding airway lumens and also in alveolar regions (Fig. 4G and fig. S3). Airway mucus occlusion was likely due primarily to high levels of mucin production and secretion (Fig. 2C-D, Fig. 3C and 3F, Fig. 4G, and fig. S3A-B), but epithelial destruction during the first two weeks after infection, as observed previously (23, 30, 31), with a failure to restore a normal mucociliary epithelial mosaic during the following five weeks (Fig. 5 and fig. S5) probably contributed as well. Other possible mechanisms contributing to airway mucus occlusion might include airway surface liquid depletion due to inflammation during viral infection that has been observed by others but was not examined here (42). The presence of mucus in alveolar regions (Fig. 4G and fig. S3A) was more likely due to mucin synthesis and secretion by bronchiolized epithelium than by reflux from the conducting airways because of the tapering anatomy of airways and inability of terminal bronchioles to produce mucin (32, 43), the presence of intraepithelial mucin in bronchiolized alveolar epithelium indicating local production (fig. S3A), and the greater abundance of Muc5b relative to Muc5ac in alveolar regions compared to conducting airways suggesting different sources of mucus (fig. S3A). Both the number of macroscopic lung nodules (Fig. 4B) and the extent of airway mucus occlusion (Fig. 4H) were markedly reduced by treatment with O/P.

Our study has several limitations in predicting whether O/P will be an effective therapeutic agent in humans. These include the inherent differences between mice and humans in anatomy, molecular biology, and interactions with pathogens. Offsetting this limitation, SeV is a paramyxovirus closely related to respiratory syncytial virus and parainfluenza viruses that cause considerable respiratory disease in humans, and SeV is a natural pathogen of rodents that replicates in their lungs and causes injury resembling that of human paramyxoviral infections (6, 8, 11, 23, 30, 31). In addition, the efficacy of O/P in inducing epithelial resistance to viral infection in human airway cells has been demonstrated in vitro (21). Another limitation is that SeV infection of the lungs was induced in mice by bolus aspiration, whereas human infection most commonly proceeds by inhalation of small infected droplets from the upper respiratory tract. However, this should mostly affect the timing of treatment, which can be determined empirically in human subjects, rather than the underlying therapeutic principle.

Treatment of susceptible human subjects with O/P has the potential to prevent the substantial morbidity associated with respiratory viruses in the pathogenesis of airway diseases including asthma, COPD, and cystic fibrosis. We have treated 49 normal human subjects in a single-dose, ascending, dose-strength trial (NCT02124278), and a multiple-dose safety and tolerability trial (NCT02566252). Treatment was well tolerated, and we believe we reached the upper plateau of the dose-response curve based upon small rises in blood neutrophils and C-reactive protein levels resembling the small sigmoidal-shaped plots of rises in blood IL-6 levels in mice (15). Next steps in development of the drug include viral challenge studies in subjects with airway disease (44-46), followed by natural infection preemption studies in subjects with airway disease or immune compromise (20, 38, 47, 48). O/P showed reductions in acute SeV lung burden and chronic asthma phenotypes comparable to those seen with aerosolized IFN-β in mice (Fig. 8), and in a clinical trial, IFN-β administered within 24 hours of the onset of cold symptoms reduced asthma symptoms and the fall in peak expiratory flow in difficult-to-treat asthmatics (38). Together, O/P and IFN-β show additive or synergistic activity in mice (Fig. 8D and fig. S6C), which is not surprising in view of their differing mechanisms of action with O/P causing no induction of Type I interferons (12-14, 21). Whether either or both of these drugs, or some other stimulant of innate epithelial resistance to infection, can ameliorate the burden of virus-caused airway disease will be determined in future clinical trials of efficacy and safety.

## Materials and Methods

### Mice and chemicals

Seven week-old female C57BL/6J mice were obtained from the Jackson Laboratory (Sacramento, CA) and housed in specific pathogen-free conditions on a 12-hour light/dark cycle with free access to food and water. For euthanasia, mice were injected intraperitoneally with 2,2,2-tribromoethanol (250 mg/kg) and exsanguinated by transection of the abdominal aorta. All procedures were performed in accordance with the Institutional Animal Care and Use Committee of MD Anderson Cancer Center and the Texas A&M Institute for Biosciences and Technology. Chemicals were obtained from MilliporeSigma (St. Louis, MO) unless otherwise specified.

### Viral challenge

Sendai virus (parainfluenza type 1) was obtained from the ATCC (Manassas, VA) and expanded in primary rhesus monkey kidney cells obtained from Cell Pro Labs (Golden Valley, MN). In most experiments, mice were infected with 2.4 × 10^6^ plaque forming units (pfu) in 40 µl PBS instilled into the oropharynx and aspirated into the lungs, with mice suspended by the upper incisors on a board at 60° from horizontal under isoflurane inhalation anesthesia. The dose of virus was chosen to result in severe bronchopneumonia, but with death in <20% of mice. Lower and higher concentrations of SeV were given in pilot experiments to establish an optimal dose for chronic studies (Fig. 1C), and in a high-dose challenge study (1.2 × 10^8^ pfu) to examine additive effects of O/P and IFN-β (Fig. 8D). Lung virus burden was measured by reverse transcription quantitative PCR (RT-qPCR) of the Sendai matrix (M) protein normalized to mouse 18S rRNA as described (39), except for substitution of the primers to ACTGGGACCCTATCTAAGACAT and TAGTAGCGGAAATCACGAGG, and analyzed by the comparative C_T_ method (49).

### Treatment with aerosolized O/P

This was performed as described (15). Briefly, ODN 5’ TCG TCG TCG TTC GAA CGA CGT TGA T 3’ as the sodium salt on a phosphorothioate backbone (ODN M362) was purchased from TriLink BioTechnologies (San Diego, CA), and 2,3-bis(palmitoyloxy)-2-propyl-Cys-Ser-Lys-Lys-Lys-Lys-OH (Pam2CSK4) as the trifluoroacetic acid salt was purchased from Peptides International (Louisville, KY). A solution of ODN (1 µM) and Pam2CSK4 (4 µM) in endotoxin-free sterile water (8 ml) was placed in an Aerotech II nebulizer (Biodex Medical Systems, Shirley, NY) driven by 10 l/min of 5% CO_2_ in air to promote deep breathing. The nebulizer was connected by polyethylene tubing (30 cm × 22 mm) to a 10 l polyethylene chamber vented to a biosafety hood. Mice were exposed to the aerosol for 20 min, resulting in the nebulization of ∼4 ml of O/P solution. In one set of experiments determining dose-response relationships, the concentration of O/P solution was varied as indicated.

### Lung lavage

This was performed by instilling and collecting two 1 ml aliquots of PBS through a 20 gauge cannula inserted through rings of the exposed trachea of euthanized animals, then combining the aliquots as described (15). Total leukocyte count was determined using a hemocytometer, and differential counts by cytocentrifugation of 200 μl of lavage fluid at 300 *g* for 5 min followed by Wright-Giemsa staining.

### Epithelial mucin content and lumenal mucus occlusion

Epithelial mucin content was measured as described (32, 50, 51). Briefly, lungs were fixed by intratracheal inflation with 10% formalin to 20 cm H_2_O pressure for 12 h, and then embedded in paraffin. Tissue blocks were cut into 5-µm sections, mounted on frosted glass slides (Medline, Northfield, IL), deparaffinized with xylene, washed with ethanol, then rehydrated and stained with periodic acid fluorescent Schiff reagent (PAFS). Images were acquired by investigators blinded to mouse treatment, and morphometric analysis of the images for quantitation of intracellular mucin was performed using ImagePro (Media Cybernetics, Bethesda, MD). Data are presented as epithelial mucin volume density, signifying the measured volume of mucin overlying a unit area of epithelial basal lamina. Airway mucus plugging was measured by modifications, as follows, of a method we have described previously (52). Lungs were fixed by immersion, to avoid displacement of lumenal mucus by inflation, in methanol-based Carnoy’s solution (methacarn), to minimize changes in mucus volume, for 48 h at 4°C. Lungs were then excised, and the left lung was placed in a precision cutting tool to generate 1 mm transverse slices (fig. S3D). Four slices were embedded in paraffin, with two slices taken below the hilum and two above, each separated by a 1 mm slice that was discarded. One 5 µm section was taken from each slice, deparaffinized, rehydrated, and then stained with PAFS. The total area of lumenal mucus in all 4 slices was measured for each lung using ImageJ software (53).

### Lung mechanics

These were analyzed using a flexiVent (Scireq, Montreal, Canada) forced oscillation ventilator system as described (54). Briefly, mice were anaesthetized with urethane (2 mg/g i.p.) and paralyzed with succinylcholine chloride (500 µg i.p. followed by continuous i.p. infusion at 10 μg/g·min). They were then tracheostomized and ventilated at 150 breaths/min, 10 μl/g, against 2-3 cm H_2_O positive end-expiratory pressure. Respiratory resistance was assessed at baseline and in response to three incremental doses of aerosolized methacholine (MCh) (3, 10 and 30 mg/ml) administered by an in-line ultrasonic nebulizer (4-6 μm, Aerogen, Ireland). Total respiratory resistance was calculated by averaging five values measured for each dose of MCh for each mouse.

### Histochemistry

Cellular elements and tissue morphology were examined with hematoxylin and eosin (H&E) staining, and collagen and lipids in nodules were visualized with picrosirius red (PSR) and Sudan black B staining, respectively. For H&E and PSR staining, lungs were fixed and sectioned as above. For H&E staining, slides were incubated in Harris’ hematoxylin, eosin and graded ethanol. For PSR staining, slides were incubated in direct red 80 with picric acid for 1 h, then rinsed with acidified water. To image lipids, fixed lungs were cut into lobes and immersed in 10% Optimal Cutting Temperature compound (Tissue-Tek OCT; Sakura, Torrance, CA) and 20% sucrose in PBS on a rocker at 4° C overnight. Tissue was then frozen in OCT, cut into 10-µm sections, and mounted on Superfrost Plus glass slides (Fisher Scientific, Hampton, NH) for this and all other studies using frozen tissue. Sections were stained with Sudan black B (Rowley Biochemical, Danvers, MA) for 1 h and counterstained with Kernechtrot’s nuclear fast red (Rowley) for 20 min, followed by mounting with AquaMount medium (Polysciences, Warrington, PA). All images were acquired with an Olympus BX 60 microscope at 10x or 40x objectives unless otherwise specified.

### Immunohistochemistry

Mouse lungs were fixed, embedded, sectioned, and deparaffinized as above, then exposed for 10 min to 3% H_2_O_2_ in 90% methanol, and then heated for 10 min in 10 mM sodium citrate, pH 6.0, for antigen retrieval. Slides were rinsed in water, blocked in horse serum (Vector Laboratories, Burlingame, CA) or goat serum (Jackson ImmunoResearch, West Grove, PA) for 1h, then rinsed again and incubated with primary antibodies diluted in blocking solution at 4° C overnight. Primary antibodies used were: goat anti-mouse IL-33 (AF3626, R&D Systems, Minneapolis, MN; 1:1000), rabbit anti-Muc5ac (gift from Dr. Camille Ehre, 1:500) (55), and mouse monoclonal antibody MDA-3E1 (1:500) to detect Muc5b (54). After incubation, secondary antibodies – biotinylated horse anti-goat IgG (Vector), horseradish peroxidase (HRP)-labelled goat anti-rabbit (Millipore-Sigma) or HRP-labelled goat anti-mouse antibody (ThermoFisher Scientific) – were added for 2 h at room temperature. Tissue sections were then washed with PBS, counterstained with H&E, and mounted with VectaMount (Vector).

### Immunofluorescence

Lungs were frozen in OCT and cut into 10-µm sections as above. Sections mounted on glass slides were thawed, washed with water, blocked with donkey serum (Jackson ImmunoResearch), then incubated with primary antibodies overnight at 4°C. Primary antibodies used were: rabbit anti-prosurfactant protein C (AB3786, EMD Millipore, Burlington, MA; 1:1000), rabbit anti-keratin 14 (KRT14, RB-9020-P0, Thermo Fisher, Waltham, MA; 1:500), rat anti-mouse CD68 (FA-11, BioLegend, San Diego; 1:200), and anti-Muc5ac and anti-Muc5b as above. This was followed by the addition of secondary antibodies conjugated to Alexa 555 (Cy3), Alexa 488 (FITC) and DAPI (Jackson ImmunoResearch) for 2 h. A confocal microscope (A1plus, Nikon) was used to acquire all images. For quantitative studies, random images were acquired by investigators blinded to subject identity. Percentages of IL-33 positive cells were counted using ImageJ (Fig. 6C-E). To measure IL-33 fluorescence intensity (Fig. 6B), the left lung was sectioned at the axial bronchus between lateral branches 1 and 2 using a precision cutting tool (fig. S4), then imaged using an upright microscope (Olympus BX 60) with a 40x lens objective and identical parameters of exposure time, color intensity, contrast and magnification. Images were uploaded to ImageJ, and a red (IL-33) background intensity threshold was established by first measuring fluorescence intensity for background regions (those without blue nuclear staining) and regions of interest (double-positive red and blue staining). The threshold was then set as the mean between maximal background regions and mean regions of interest, and subtracted from all fields. The “Oval” tool was used to identify IL-33+ nuclei, these were imaged in grey scale (Fig. 6B inset), and grey scale values for all pixels in a field were plotted as arbitrary fluorescence intensity units (Fig. 6B).

### Whole mount immunofluorescence

Fixed lung lobes were embedded in OCT as above, then frozen sections of 100∼200 µm thickness were cut. Sections were thawed and blocked in 2 ml Eppendorf tubes containing PBS with 0.3% TritonX-100 and 2.5% donkey serum (017-000-121, Jackson ImmunoResearch) on a rocker at room temperature for 2 h. After blocking, lung tissues were incubated with primary antibodies diluted in PBS with 0.3% TritonX-100 on a rocker at 4° C overnight. Primary antibodies used (Fig. 5) were: rat anti-E-cadherin (E-Cad, 131900, Invitrogen; 1:1000), goat anti-CCSP (1:2,500, a gift from Dr. Barry Stripp), and mouse anti-acetylated tubulin (Ac-Tub, T6793, MilliporeSigma; 1:2,000). On the second day, lung tissues were washed in PBS with 1% TritonX-100 and 1% Tween-20 for 3 h, then incubated with secondary antibodies diluted in PBS with 0.3% Triton X-100 on a rocker at 4° C overnight. Fluorescent secondary antibodies were obtained from Jackson ImmunoResearch and used at 1:1,000. On the third day, lung tissues were again washed in PBS with 1% TritonX-100 and 1% Tween-20 for 3 h, washed with PBS twice for 5 min, fixed with 2% PFA for 2 h on a rocker, and washed with PBS for 1 h. After the PBS wash, lung tissues were mounted with AquaMount and airways of the left main bronchi were imaged on a Nikon A1plus confocal microscope. For each lung, six image fields – each consisting of several hundred epithelial cells – were counted.

### Treatment with aerosolized IFN-β

In time-course and dose-finding pilot studies, mice were treated with varying concentrations of aerosolized IFN-β (R&D Systems, Minneapolis, MN) with 0.05% BSA in water (8 ml) as described above for O/P, except that the aerosolization was continued until all IFN-β was delivered (approximately 40 min). At varying times after treatment, mice were euthanized and underwent bronchoalveolar lavage for measurement of cytokines. In efficacy studies, mice were treated with 400,000 units of aerosolized IFN-β, then challenged the next day with Sendai virus by oropharyngeal instillation and euthanized five days later for measure of lung virus burden.

### Statistical analyses

All data sets were first analyzed with the Shapiro-Wilk test to determine normality. For analyses where there was only a single comparison between two groups, data was analyzed by Student’s *t*-test or Mann-Whitney *U* test for normally and non-normally distributed data, respectively. For analyses where multiple experimental groups were compared against a single control, data were first analyzed by ANOVA on ranks to determine if a significant difference between any groups was present. If found, data were further analyzed using Dunn’s test and significance determined from adjusted *P* values. For analyses where each experimental group was compared to every other group, data were first analyzed by one-way ANOVA or ANOVA on ranks for normally and non-normally distributed data, respectively. If a significant difference between any groups was present by ANOVA, data were further analyzed using Holm-Sidak’s test for one-way ANOVA or Dunn’s test for all pairwise combinations for ANOVA on ranks. Significance was determined from adjusted *P* values. For studies comparing O/P and IFN-β, an interaction score from linear regression analysis was used to determine whether there was a synergistic effect of the combination compared to either drug alone. All data were analyzed using SigmaPlot (version 12.5, Systat Software). *P* < 0.05 was considered statistically significant.

## Supplementary Materials

Fig. S1. Lung SeV burden days 3 through 7 after infection.

Fig. S2. Effect of O/P treatment 20 days before or after SeV infection.

Fig. S3. Staining for Muc5ac and Muc5b, collagen, and macrophages in lung nodules.

Fig. S4. Location of left axial bronchus sections for mucin content, and image of the precision cutting instrument.

Fig. S5. Whole mount and cross-sectional immunofluorescence staining of airway epithelium 14 days after Sendai infection.

Fig. S6. Lung lavage fluid cytokine responses to aerosolized O/P and IFN-β.

## Acknowledgements

The authors thank Mark McArthur, DVM, for helpful discussions of pathologic findings.

## Author contributions

BLS, SEE, MJT, BFD conceived the study; DLG, JRF, GV, AMJ, ST, SW, BH, JC performed the studies; DEO oversaw the statistical analyses; MJT made the figures; BFD wrote the manuscript.

