## supplement for "Aerosolized TLR Agonists Suppress Acute Sendai Virus Lung Infection and Chronic Airway Disease in Mice"

**Supplementary Materials**

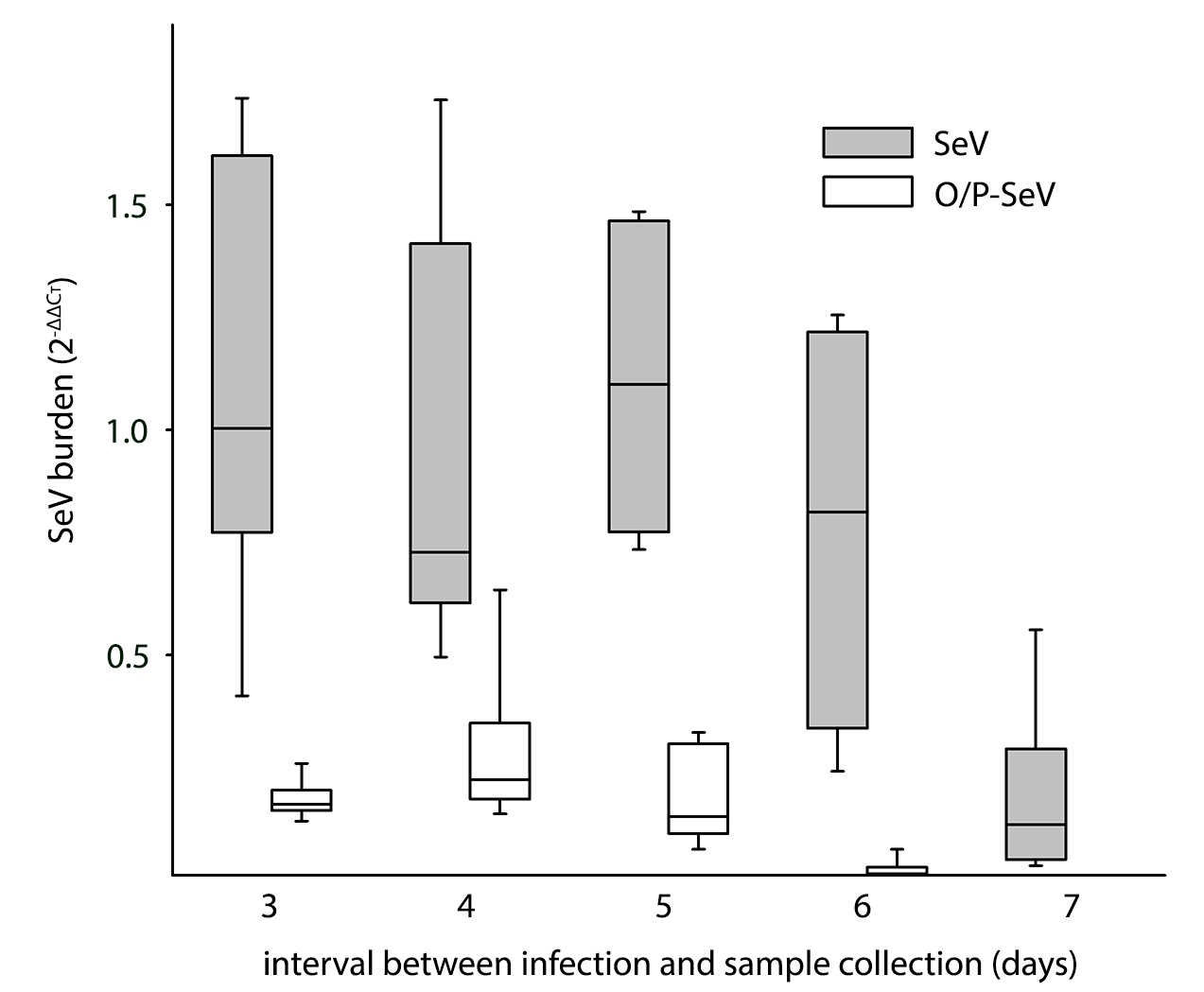

**Fig. S1.** **Lung SeV burden on days 3 through 7 after infection**. Mice were treated or not with O/P one day before infection with SeV, then lung SeV burden was measured by qRT-PCR at various days after infection. For mice treated with O/P, there was a significant reduction in SeV burden compared to untreated mice on all days. (Boxes show median and interquartile range, whiskers show 10^th^ and 90^th^ percentiles; N = 6 mice/group.) There was no significant difference in SeV burden in untreated mice at days 3, 4 or 6 compared to day 5, but there was a significant reduction in SeV burden on day 7 compared to all other days (*P<*0.05 for one-way ANOVA with Holm-Sidak’s test for multiple comparisons, asterisk not shown).

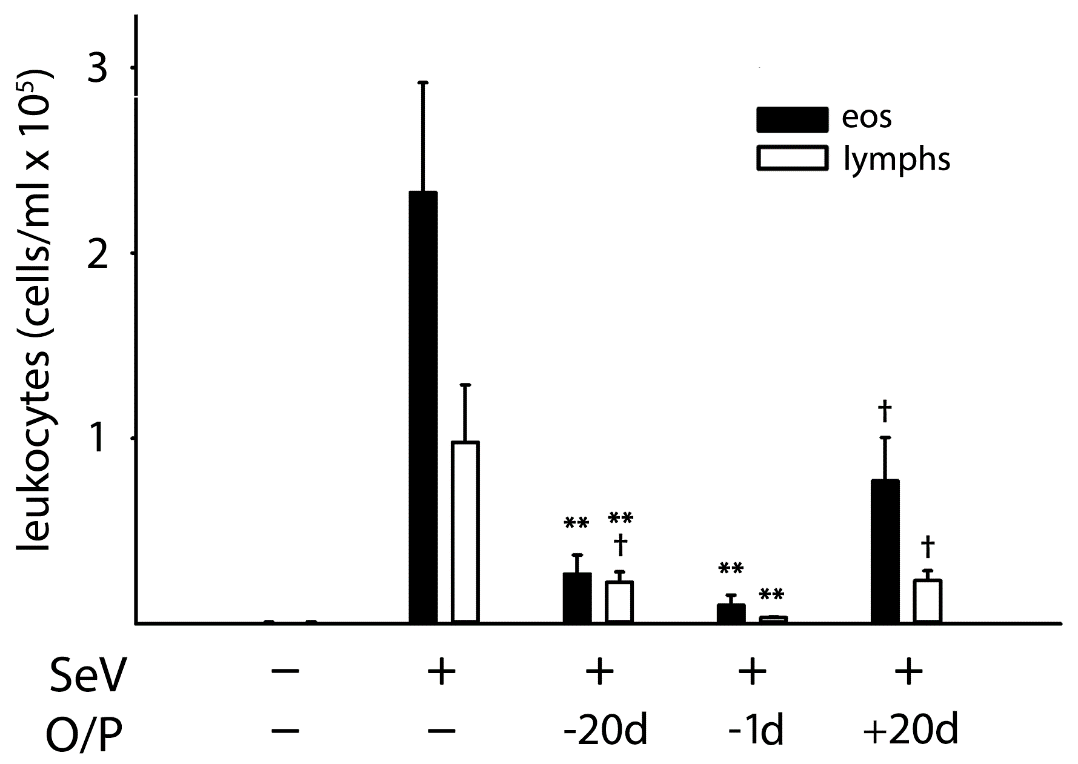

**Fig. S2. Effect of O/P treatment 20 days before or after SeV infection on lung leukocytes 49 days after infection.** Mice were treated with O/P outside of the time interval during which O/P has been shown to have an antiviral effect (8 days before through 3 days after infection), and compared to mice not treated with O/P and to mice treated with O/P during an interval of maximal activity (1 day before infection). Eosinophils and lymphocytes in lung lavage fluid were enumerated 49 days after SeV infection. (Bars show mean ± SEM; ***P* < 0.01 for [SeV^+^,O/P^+^] *vs* [SeV^+^,O/P^-^], and ^†^*P* < 0.05 for mice treated 20 days before or after SeV infection compared to those treated 1 day before infection, by ANOVA on ranks with Dunn’s test for multiple comparisons; N = 13-18 mice/group pooled from 3 experiments.)

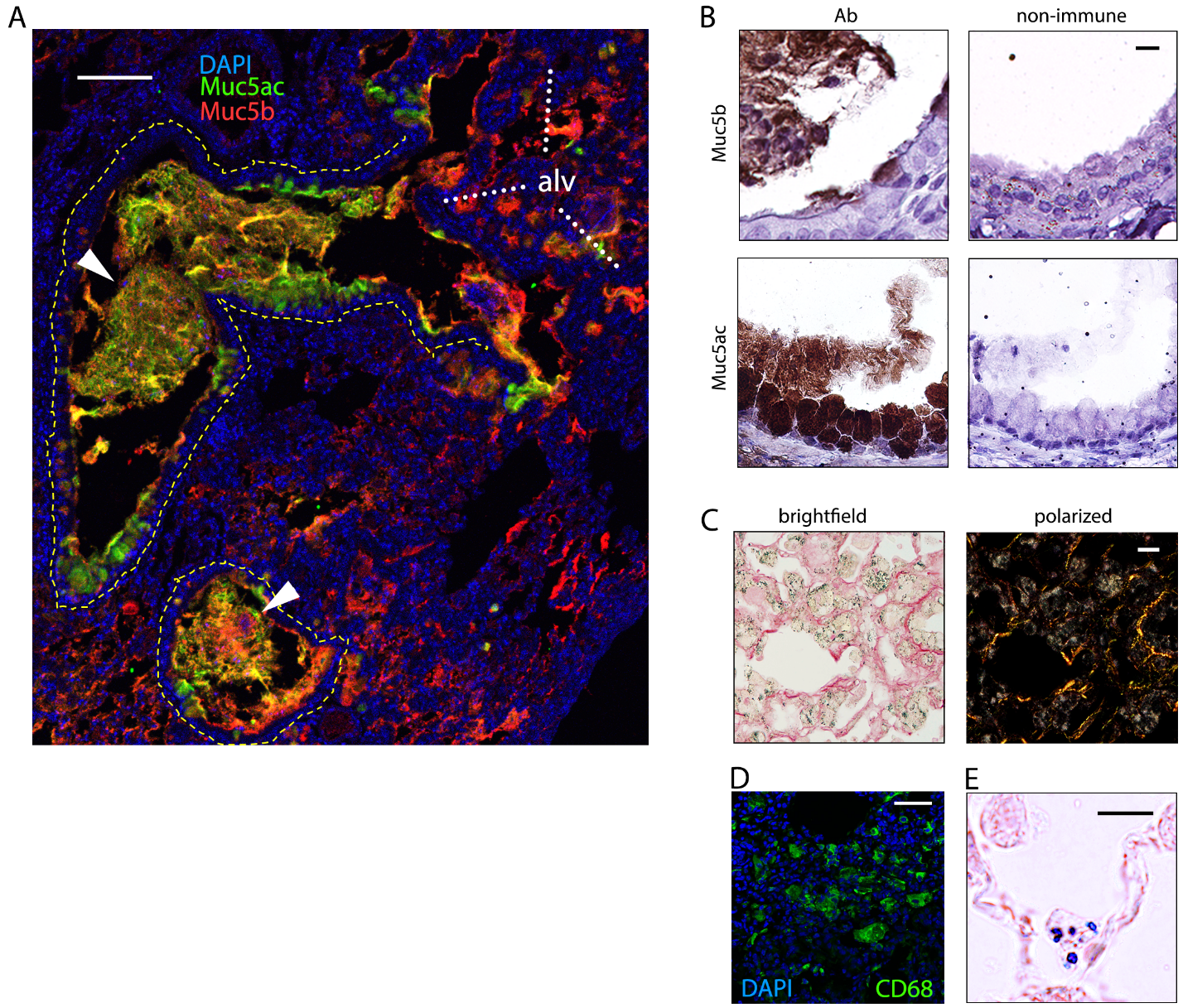

Fig. S3. (A) Staining for Muc5ac and Muc5b, collagen, and macrophages in lung nodules 49 days after SeV infection. Section through a lung nodule showing immunofluorescence staining for the secreted airway mucins Muc5ac and Muc5b, and DAPI staining for nuclei. White arrowheads indicate airway lumenal mucus plugs, and the epithelium of both airways is outlined with white dashed lines. An alveolar region is indicated by white letters “alv” and heavy white dotted lines. Scale bar = 100 µm. (B) Immunohistochemical staining of airways plugged with mucus for Muc5b and Muc5ac. Scale bar = 10 µm. (C) Brightfield and polarized microscopy of the same section of a nodule stained with picrosirius red. Scale bar = 20 µm. (D) Immunofluorescence staining for CD68 to confirm the identity of numerous macrophages in alveolar spaces. Scale bar = 50 µm. (E) Sudan black staining of an alveolar T2 cell in a healthy lung for comparison to staining of macrophages in lung nodules in Figure 4(F). Scale bar = 10 µm.

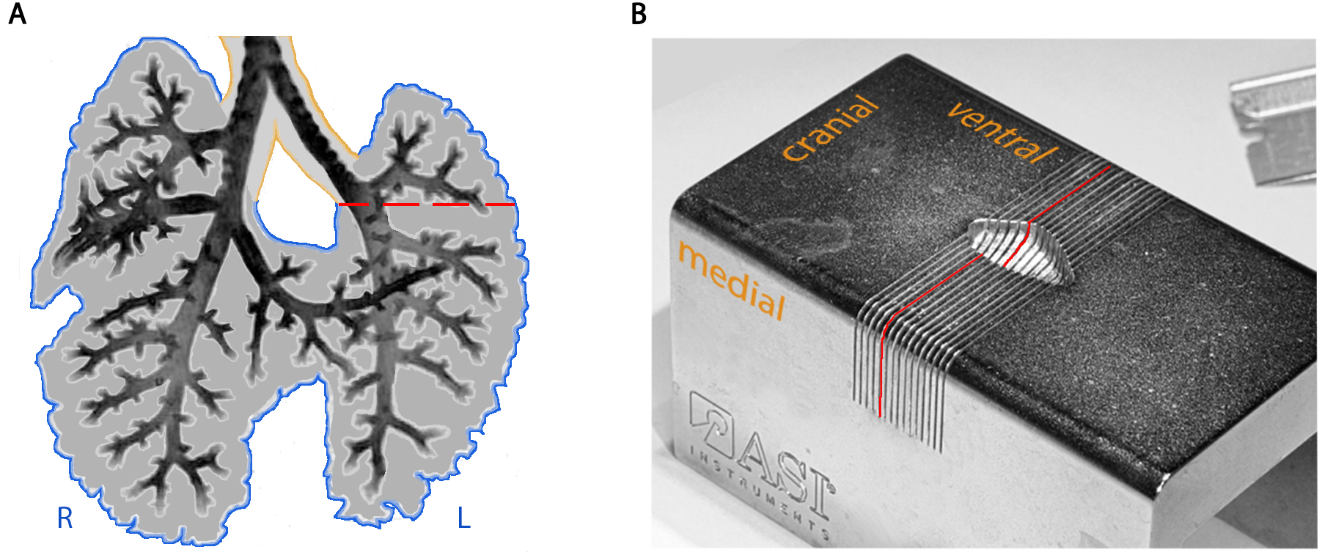

Fig. S4. Location of left axial bronchus sections to assess mucin content, and photograph of the precision cutting instrument. (A) Schematic of mouse lung airway anatomy with a dashed red line showing the location of initial cut, with two 1 mm slices then taken above this plane and two slices below. R indicates right lung, and L indicates left lung. Image shows day E15 lungs stained immunohistochemically for Sox2 to demonstrate airways, modified from Alanis DM, et al, *Nat Commun* 2014, 5:3923. (B) The precision cutting instrument used to generate the 1 mm lung slices, with a red line corresponding to the cut shown in A, and the orange lettering indicating the placement of the left lung into the device in relation to its cranial, ventral and medial surfaces.

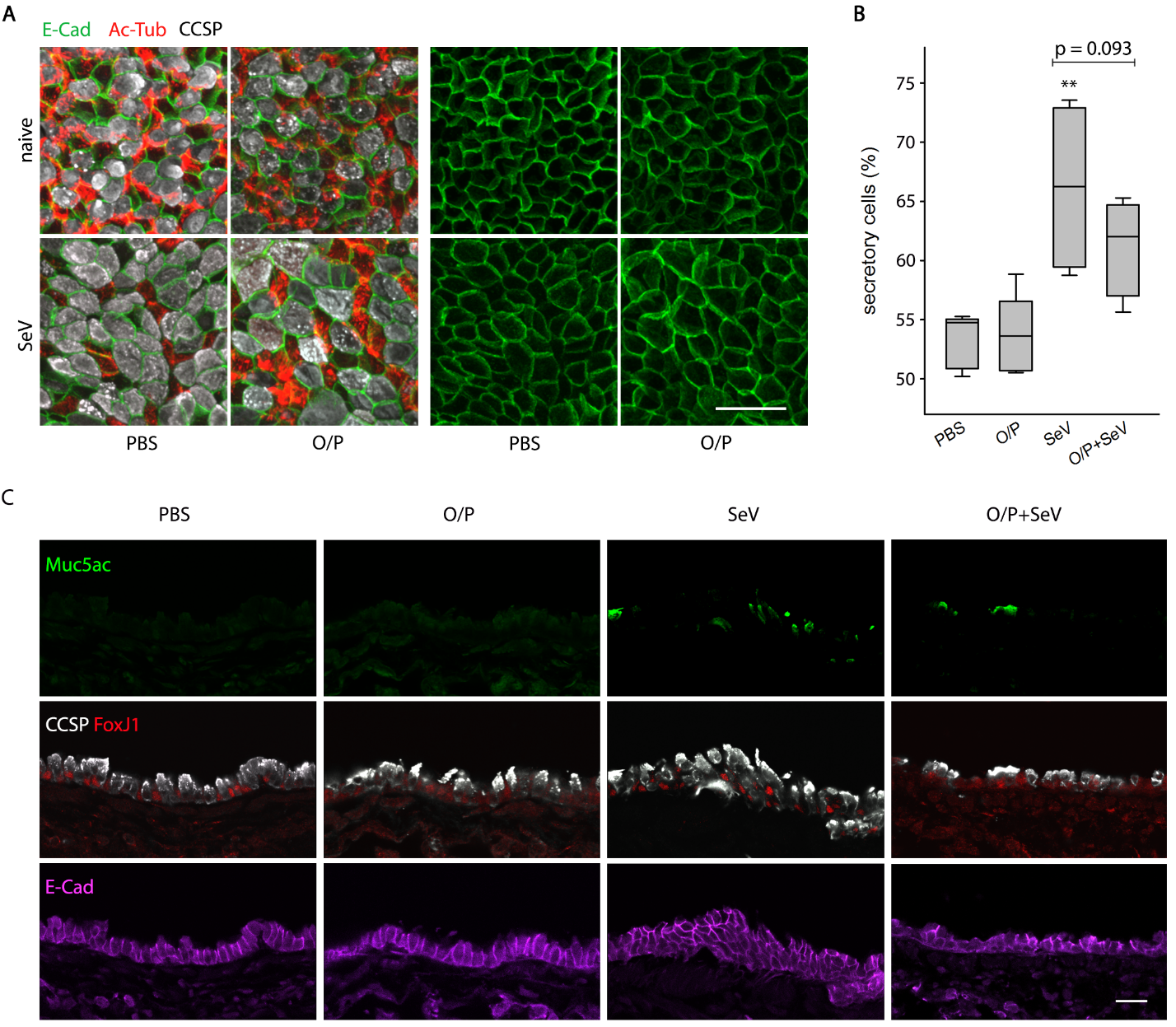

**Fig. S5. Whole mount and cross-sectional immunofluorescence staining of airway epithelium 14 days after SeV infection.** **(A)** Whole mount immunofluorescence images of the axial bronchi of mice, treated or not with O/P, then infected or not with SeV, 14 days earlier. The four images on the left show immunofluorescence staining for E-cadherin (E-Cad), acetylated tubulin (Ac-Tub), and club cell secretory protein (CCSP) to identify changes in numbers and sizes of individual cell types, whereas the four images on the right show staining only for E-cadherin to best illustrate changes in cell shape and pattern. Scale bar = 20 µm. (**B**) Quantification of the percentage of CCSP-positive secretory cells in images such as those in A. (Boxes show median and interquartile range, whiskers show 10^th^ and 90^th^ percentiles; ***P*<0.01 by Student’s *t*-test for [SeV^+^,O/P^-^] *vs* [SeV^-^,O/P^-^]; N = 5 mice/group with 6 images/mouse, all acquired from the axial bronchus, 3 at L2-3 and 3 at L4-5.) (**C**) Cross-sectional images of the same airways as in (A) showing immunofluorescence staining for Muc5ac (top row), CCSP to identify secretory cells and FoxJ1 to identify ciliated cells (middle row), and E-cadherin to illustrate epithelial thickness and cell outlines (bottom row). Scale bar = 20 µm.

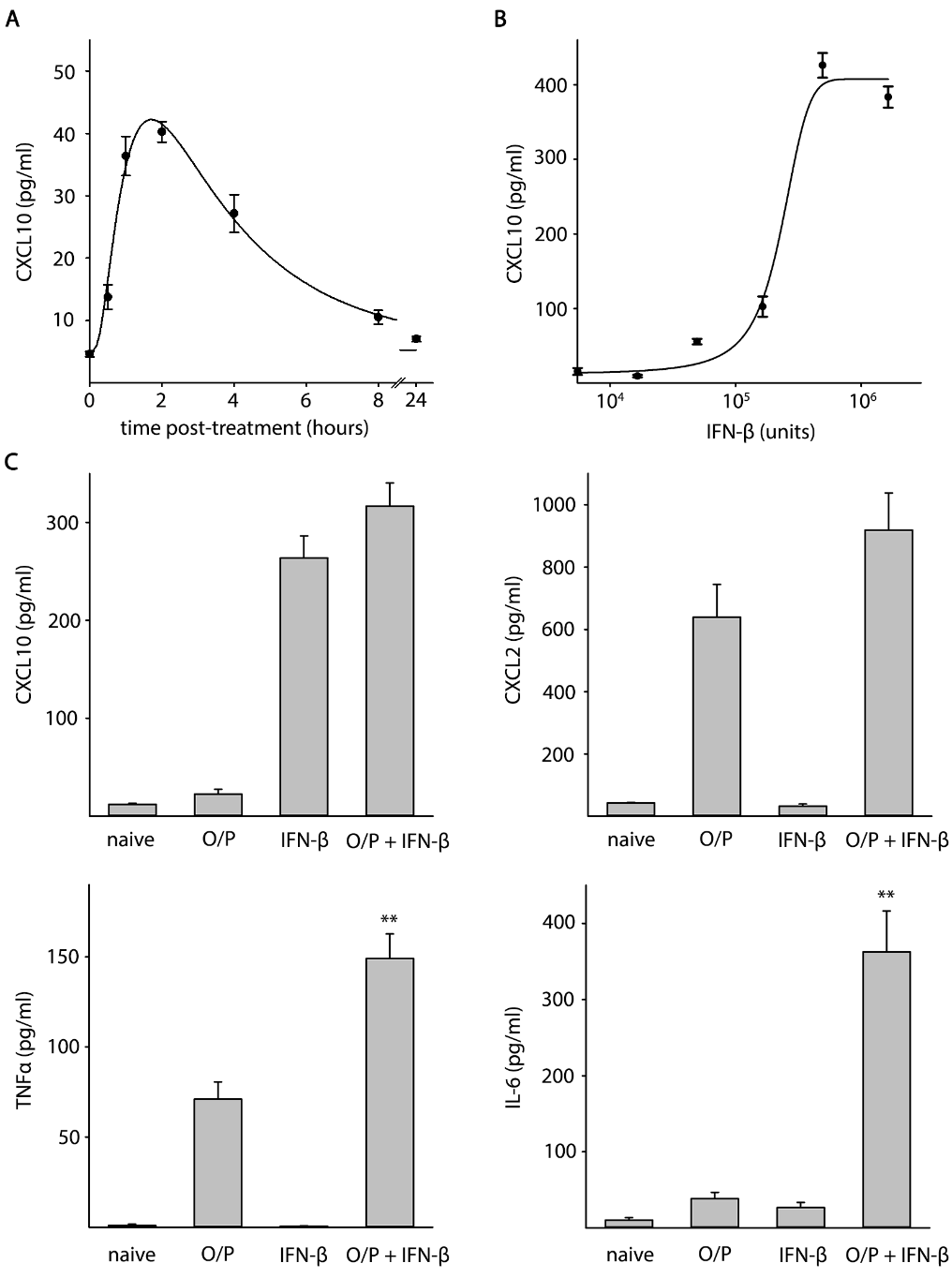

**Fig. S6. Lung lavage fluid cytokine responses to aerosolized O/P and IFN-β.** (**A**) IFN-β (205,000 units) was administered by aerosol to a cohort of mice, and mice in groups of 4-6 were sacrificed at the indicated time points for measurement of CXCL10 in lung lavage fluid. (Plotted points show the mean ± SEM.) (**B**) Mice in groups of 5 were administered varying amounts of IFN-β by aerosol, then were sacrificed after 2 h for measurement of CXCL10 in lung lavage fluid. (Plotted points show the mean ± SEM.) (**C**) Mice in groups of 5 were treated or not with O/P aerosolized 24 h before analysis, or with 400,000 units of IFN-β aerosolized 2 h before analysis, or with both drugs, and the indicated cytokines were measured in lung lavage fluid. (Bars show mean ± SEM; ***P* < 0.01 for a synergistic interaction between O/P and IFN-β by linear regression for the interaction value between the two drugs compared to either drug alone; N = 4 mice/group).
